# Silent dissemination of plasmid-borne tigecycline resistance gene *tet*(X6) in livestock-associated *Acinetobacter towneri*

**DOI:** 10.1101/2020.05.08.085514

**Authors:** Yingying Cheng, Yong Chen, Yang Liu, Jingjie Song, Yuqi Guo, Yanzi Zhou, Tingting Xiao, Shuntian Zhang, Hao Xu, Yunbo Chen, Tongling Shan, Yonghong Xiao, Kai Zhou

**Author notes:** These authors contributed equally to this work (Yingying Cheng, Yong Chen). Corresponding author. Mailing address: Dongmen North Road No. 1017, Shenzhen People’s Hospital, 518020 Shenzhen, China.

## Abstract

A tigecycline-resistance gene, *tet*(X6), was detected on a 159-kb novel plasmid carried by a tigecycline-susceptible livestock-associated *Acinetobacter towneri* isolate. The genetic context of *tet*(X6) (ΔIS*Vsa3*-*tet*(X6)-*abh*-*guaA*-IS*Vsa3*) is highly similar with that of the other plasmid-borne *tet*(X) variants. The 23-Ala residue of the first FAD binding site conferred higher activity to Tet(X6) than the 23-Gly reside conserved in the other plasmid-borne *tet*(X)s. To our knowledge, this is the first report of *tet*(X6) carried by the plasmid.

---

Tigecycline is one of the last resorts to treat clinical infections caused by multi-drug resistance (MDR), especially carbapenem-resistant bacteria (1). Tet(X) family contains a series of flavin-dependent monooxygenases, which can degrade tetracycline and tigecycline to 11a-hydroxy-oxytetracycline and 11a-hydroxytigecycline by a similar enzymatic modification pattern, respectively (2, 3). The first member of the Tet(X) family was identified in Tn*4351* and Tn*4400* carried by the chromosome of anaerobe *Bacteroides fragilis* (4). Recently, three plasmid-borne *tet*(X) variants mediating tigecycline resistance, *tet*(X3), *tet*(X4) and *tet*(X5), were consecutively detected in *Acinetobacter* and *Enterobacterales* isolates obtained from animals, animal-derived foods, and humans in China (5-7). These findings warn the wide-dissemination possibility of tigecycline resistance in clinical setting mediated by mobile genetic elements. Most recently, *tet*(X6) was identified on the chromosome of *Proteus* spp., *Myroides* spp. and *Acinetobacter* spp. Strains (8, 9), and on integrative and conjugative elements ICE*Pgs6Chn1* carried by the chromosome of *Proteus* spp. (9, 10). We here first report the detection of a plasmid-borne *tet*(X6) in a livestock-associated *A. towneri* strain.

PCR was performed with use of universal primers *tet*(X)-F (5’-TGCTTGAACCTGGTAAGAAG-3’) and *tet*(X)-R (5’-AATGAGCAGCATCGCCAATC-3’) to screen *tet*(X) variants in 290 *Acinetobacter* spp. strains isolated from livestock stool samples collected in China in 2019. A *tet*(X6)-positive *A. towneri* strain AT205 recovered from a swine fecal sample was detected. AT205 exhibited resistance to gentamicin and some tetracycline families, but susceptible to tigecycline with the minimum inhibitory concentration (MIC) at 1 mg/L (Table S1).

To understand the vector of *tet*(X6), AT205 was sequenced by using Hiseq 4000 system (Illumina, San Diego, US) and PromethION platform (Nanopore, Oxford, UK). Hybrid assembly using Unicycler version 0.4.8 (11) resulted in a 2.66-Mb circular chromosome (confirmed by PCR) with GC content of 41.48% (CP048014), a completed plasmid pAT205 (confirmed by PCR) with a size of 158.867 kb (CP048015) and three contigs (CP048016-CP048018). The *tet*(X6) gene was located on the plasmid pAT205 (Figure 1A). This is different with the recent reports that *tet*(X6) was carried on the chromosome of *Proteus* spp., *Myroides* spp. and *Acinetobacter* spp. strains (8-10). Therefore, this is the fourth plasmid-borne *tet*(X) variant identified after *tet*(X3), *tet*(X4) and *tet*(X5) (6, 7). Notably, *Acinetobacter* spp. is the common host of *tet*(X3), *tet*(X5) and *tet*(X6) (6-8), suggesting that the genus may be an important reservoir for the plasmid-borne tigecycline-resistance genes.

**Figure 1.**
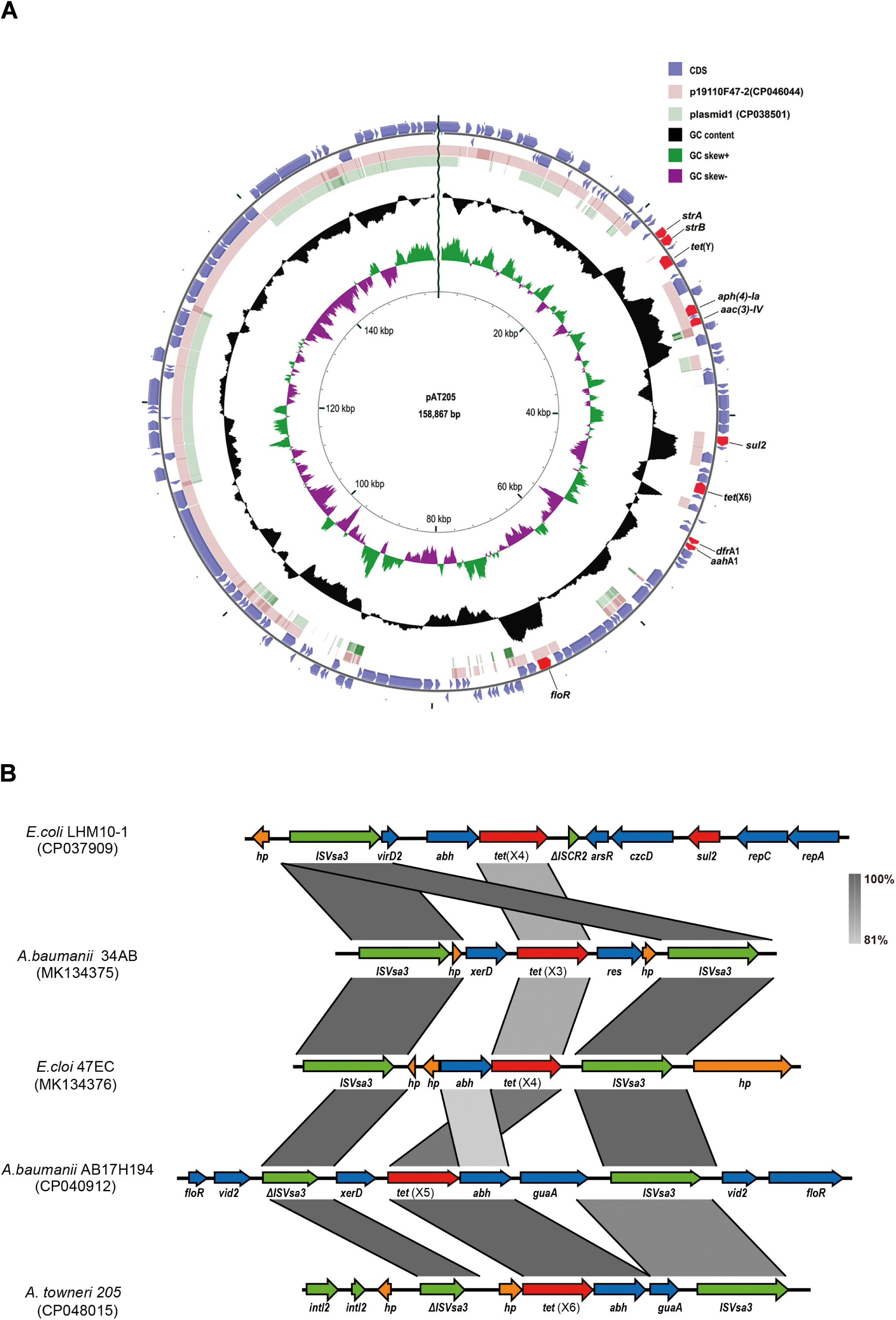
The structure of *tet*(X6)-carrying pAT205. (A) The circles from outside to inside depict: (i) the antibiotic resistance genes harbored by pAT205 are marked in red; (ii) the positions of predicted coding sequences transcribed in clockwise and counterclockwise orientation in indigo blue; (iii) reported homologe plasmid of pAT205 with coverage > 35% in pink and light green; (iv) the GC content plotted against 50%; (v) GC skew [(G-C)/(G+C)] in a 10000 bp window; (vi) the backbone of pAT205 with size scale in bp. The CGView Server (13) was used to generate the blast results. (B) Comparison of genetic context of plasmid-borne *tet*(X) variants. Sequence comparison and map generation were performed by using Easyfig 2.2.3 (14). Arrows represent the genes with different directions. Regions of >80% homology are shown by grey shading. Genes are colour-coded, depending on functional annotations: red, antimicrobial resistance; green, mobile elements; blue, other functions; orange, hypothetical proteins.

The genetic context of *tet*(X6) detected on pAT205 was “ΔIS*Vsa3*-*tet*(X6)-*abh*-*guaA*-IS*Vsa3*”. This is highly similar with that of plasmid-borne *tet*(X3), *tet*(X4) and *tet*(X5) previously reported (5-7) (Figure 1B). It has been shown that the IS*Vsa3*-flanking region is able to become a circular intermediate facilitating the transfer of *tet*(X3) and *tet*(X4) (8). Given the similar structure, we suppose that *tet*(X6) could be mobilized through this unit. pAT205 failed to be transferred to *A. baumannii* ATCC17978 and *E. coli* EC600 by the conjugation assay, which is due to the lack of functional conjugative system on pAT205 (only one type IV secretory gene was detected).

Blasting the sequence of pAT205 showed only two plasmids with coverage >35% (Figure 1A); p19110F47-2 (CP046044) carried by an *A. towneri* strain (99.98% identity; 66% coverage) and plasmid 1 (CP038501) carried by an *A. baumannii* strain (99.63% nucleotide identity; 35% coverage). This suggests that pAT205 is a novel plasmid, and might originate from *Acinetobacter* spp.. A class 2 integron (50498 bp-67325 bp) region carrying the cassette array of *dfrA1, sat2* and *aadA1* was identified on pAT205 (Figure S1A). The 3’ conserved segment (3’-CS) of the integron was consist of five *tns* genes (*tnsA, tnsB, tnsC, tnsD* and *tnsE*) (12), and was disrupted by a 4802-bp fragment encoding a retron-type RNA-directed DNA polymerase and IS*66* family mobile elements (Figure S1A). A Tn*6205*-like structure (22252bp-34180bp) was additionally detected on pAT205 (Figure S1B). Compared with the original structure of Tn*6205* (CP003505), 5’ and 3’ end of the Tn*6205*-like structure was reversed, and more resistance genes were carried, including a tetracycline resistance regulatory gene *tet*(R), an efflux pump gene *tet*(Y) and 2 aminoglycoside resistance genes [*aph(4)-Ia* and *aac(3)-IV*]. Such plasmid represents a high threat to public health since it confers MDR including the resistance to the last resort antibiotics.

To verify the activity of *tet*(X6), the fragment between 107 bp upstream and 29 bp downstream of *tet*(X6) including the predicted promoter was amplified using primers pUC19-*tet*(X6)-F (5’-cgctgcagGCAATTGACTTTCCGAACGG-3’) and p-*tet*(X6)-R (5’-cgtctagaTTTCTCTTTCATTTCCTCGCC-3’). The resulted product was ligated into pUC19 to construct transformant pUC19-*tet*(X6) and overexpressed in *E. coli* DH5α. In parallel, *tet*(X3) and *tet*(X4) were cloned into pUC19 as positive controls. The tigecycline MIC of the transformant DH5α-pUC19-*tet*(X6) was 8 mg/L, which was slightly lower than that of *tet*(X3) and *tet*(X4) transformants as 16 mg/L (Figure 2A). The tigecycline MIC of the negative control was 0.125 mg/L (Table S1). To compare the activity of Tet(X6) with that of Tet(X3) at the low expression level, *tet*(X6) and *tet*(X3) were cloned into pJN105 using the inducible pBAD promoter, respectively. The tigecycline MIC of DH5α-pJN105-*tet*(X6) and DH5α-pJN105-*tet*(X3) increased 16-fold and 64-fold respectively (Table S2). This further supports that the activity of Tet(X6) against tigecycline was weaker than that of Tet(X3).

**Figure 2.**
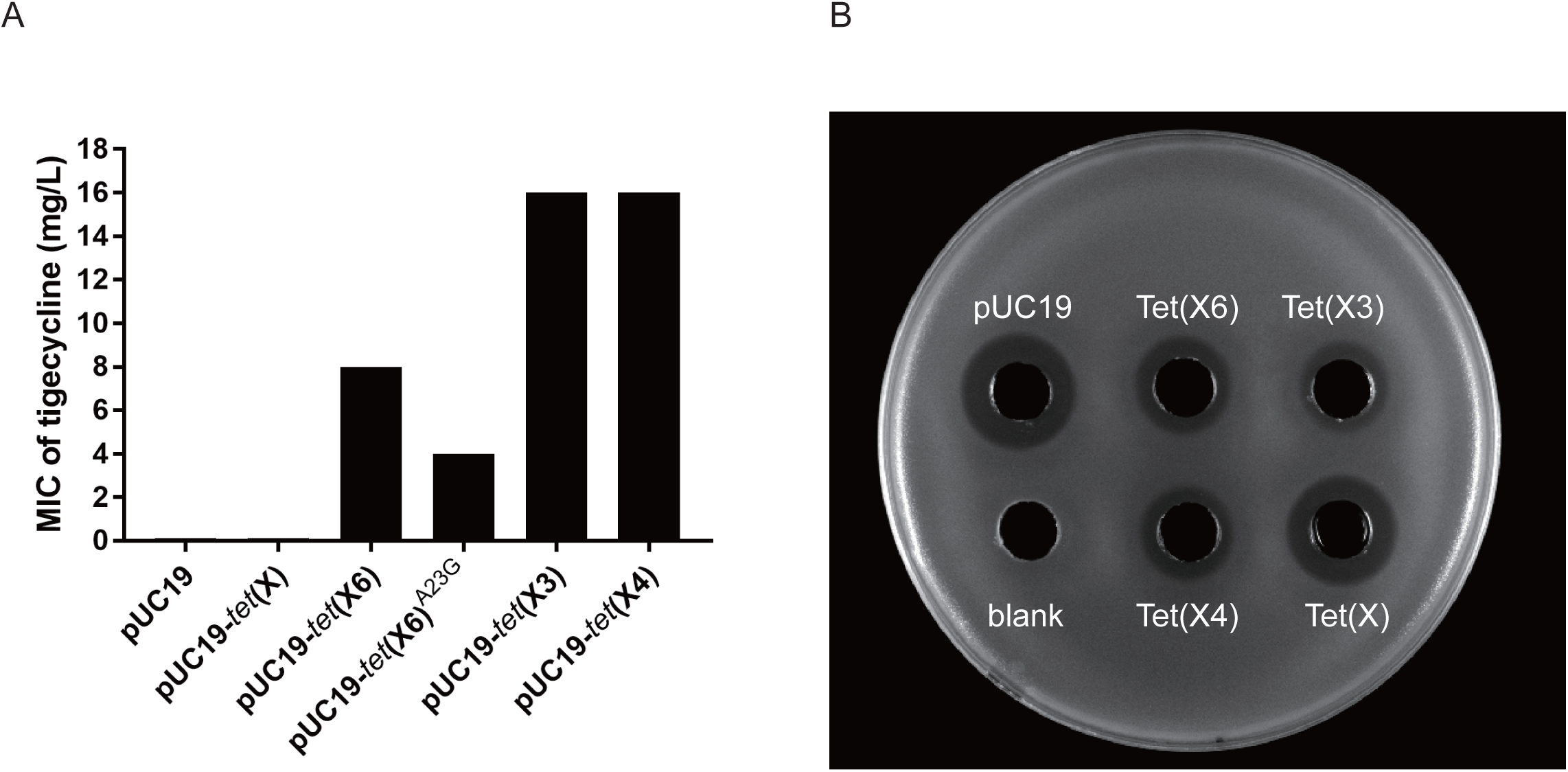
The activity of *tet*(X6). (A) The tigecycline MIC of *tet*(X) alleles overexpressed in *E. coli* DH5α. (B) The activity of Tet(X)s against tigecycline. Inhibition zones were caused by the remaining tigecycline in the supernatant of Tet(X)s-overexpressing strains, and ATCC25922 was used as the prey strain.

Microbiological degradation assay was performed as previously reported (5) with minor modification to demonstrate the tigecycline inactivation of Tet(X6). The optical density of *tet*(X)s’ clone cultures at 600 nm were adjusted to 2.0, followed by adding tigecycline into 1 mL adjusted cultures with final concentration of 2.5 mg/L. After incubation at 37°C for 8 hours, the activity of the remaining tigecycline was measured by dropping 100 μL supernatant on the LB agar plate with *E. coli* ATCC25922 as prey. The diameter of the inhibition zone caused by the supernatant of DH5α-pUC19-*tet*(X6) culture was slightly bigger than that formed by DH5α-pUC19-*tet*(X3) and DH5α-pUC19-*tet*(X4) supernatants, and smaller than that of the negative control (Figure 2B), indicating that the tigecycline inactivation capacity of Tet(X6) was slightly weaker than that of Tet(X3) and Tet(X4). This is consistent with the results described above.

We noted that the first FAD binding site of Tet(X6) was slightly different with that of the other variants in which the second Gly residue of the “GGGPVDG” motif was replaced by Ala in Tet(X6) (Figure S2). Site-directed-mutation was performed to understand whether this polymorphism has impact on the activity of Tet(X6). The left fragment of 23-Ala residue amplified using primers pUC19-*tet*(X6)-F and *tet*(X6)-SDM-AtoG-R (5’-CAACAGGCCCTCCACCAATTAT-3’) and the right fragment of 23-Ala residue amplified using primers *tet*(X6)-SDM-AtoG-F (5’-ATAATTGGTGGAGGGCCTGTTG-3’) and p-*tet*(X6)-R were fused. The fused fragment was inserted into pUC19 to yield pUC19-*tet*(X6)^A23G^. The MIC of tigecycline decreased to 4 mg/L for DH5α-pUC19-*tet*(X6)^A23G^ (Figure 2A), indicating that the 23-Ala residue of Tet(X6) was involved in the tigecycline resistance, and the site might be under selection.

In conclusion, we here first reported a plasmid-borne tigecycline-resistance gene *tet*(X6) in *A. towneri*. This is the fourth flavin-dependent monooxygenase encoding gene located on plasmid. The increasing number of plasmid-borne Tet(X)s raises a concern for the wide dissemination of tigecycline resistance in clinical setting.

## Acknowledgments

This work was supported by the National Key Research and Development Program of China (2017YFC1200200), Major Infectious Diseases Such as AIDS and Viral Hepatitis Prevention and Control Technology Major Projects (2018ZX10712-001), and the National Natural Science Foundation of China (81702045 and 81902029). We also thank Professor Jian Sun for sharing *E. coli* LHM10-1.

## Transparency declarations

None to declare.

## Data availability

The genome sequence of AT205 was submitted to GenBank under the accession numbers CP048014-CP048018.

